# Altered stomatal patterning accompanies a trichome dimorphism in a natural population of *Arabidopsis*

**DOI:** 10.1101/710889

**Authors:** Noriane M. L. Simon, Jiro Sugisaka, Mie N. Honjo, Sverre Aarseth Tunstad, George Tunna, Hiroshi Kudoh, Antony N. Dodd

## Abstract

Trichomes are large epidermal cells on the surface of leaves that are thought to deter herbivores, yet the presence of trichomes can also negatively impact plant growth and reproduction. Stomatal guard cells and trichomes have shared developmental origins, and experimental manipulation of trichome formation can lead to changes in stomatal density. The influence of trichome formation upon stomatal development in natural populations of plants is currently unknown. Here, we show that a natural population of *Arabidopsis halleri* that includes hairy (trichome-bearing) and glabrous (no trichomes) morphs has differences in stomatal density that are associated with this trichome dimorphism. We found that glabrous morphs had significantly greater stomatal density and stomatal index than hairy morphs. One interpretation is that this arises from a trade-off between the proportions of cells that have trichome and guard cell fates during leaf development. The differences in stomatal density between the two morphs might have impacts upon environmental adaptation, in addition to herbivory deterrence caused by trichome development.

## Introduction

In *Arabidopsis*, trichomes are large epidermal cells that protrude from the surface of the leaves and petioles. Trichomes play important roles in both biotic defences and abiotic stress tolerance (Levin, 1973; Mauricio and Rausher, 1997; Handley et al., 2005; Dalin et al., 2008; Sletvold et al., 2010; Sletvold and Ågren, 2012; Sato and Kudoh, 2016). However, trichome development appears to impose a fitness cost on growth and reproduction (Mauricio, 1998; Sletvold et al., 2010; Kawagoe et al., 2011; Sletvold and Ågren, 2012; Sato and Kudoh, 2016). In addition to trichomes, stomatal guard cells represent another specialized cell type that is present on the leaf surface. Trichome initiation occurs prior to stomatal meristemoid development, and the patterning of trichomes and guard cells appears to be linked (Larkin et al., 1996; Glover, 2000; Bean et al., 2002; Bird and Gray, 2003; Galdon-Armero et al., 2018). Therefore, there might be a trade-off between trichome and stomatal guard cell development during leaf formation (Glover et al., 1998).

We wished to determine whether trichome formation might be associated with changes in stomatal patterning in natural populations of plants. To achieve this, we investigated stomatal patterning in a naturally-occurring population of *Arabidopsis halleri* subsp. *gemmifera* that includes trichome-forming and glabrous morphs (Kawagoe et al., 2011; Sato and Kudoh, 2016). These trichome morph phenotypes are heritable (Sato and Kudoh, 2015, 2017). The glabrous morphs within this population harbour a large transposon-like insertion within the *GLABRA1* (*GL1*) gene (Kawagoe et al., 2011). *GL1* is also required for trichome formation in *A*. *thaliana*, with homozygous *gl1* mutants being glabrous (Oppenheimer et al., 1991). Our experiments provide new insights into the relationship between stomatal and trichome patterning under natural conditions.

## Methods

### Study site and experimental model

This investigation used a well-characterized population of *Arabidopsis halleri* subsp. *gemmifera* that is located beside a small stream in central Honshu island, Japan (Fig. 1A) (Omoide-gawa site, 35°06’ N, 134°56’ E; 230 m altitude) (Aikawa et al., 2010; Kudoh et al., 2018). *A*. *halleri* is metal tolerant and grows essentially as a monoculture at this field site because the water is contaminated by a historical mine (Kudoh et al., 2018). The species was identified by reference to herbarium and museum specimens (Kudoh et al., 2018), and a nearby population that harbours glabrous and hairy morphs supplied material for the genome sequencing and annotation of *A*. *halleri* (Briskine et al., 2017; Sato and Kudoh, 2017). The only subspecies of *A*. *halleri* present in Japan is *A*. *halleri* subsp. *gemmifera* (Honjo and Kudoh, 2019). Sampling occurred during September 2016 (photoperiod approximately 12 h, with dawn at 05:40 and dusk at 18:10). During this season, *A*. *halleri* bore larger rosette leaves that are well-suited for quantification of stomatal density (Fig. 1B).

**Figure 1.**
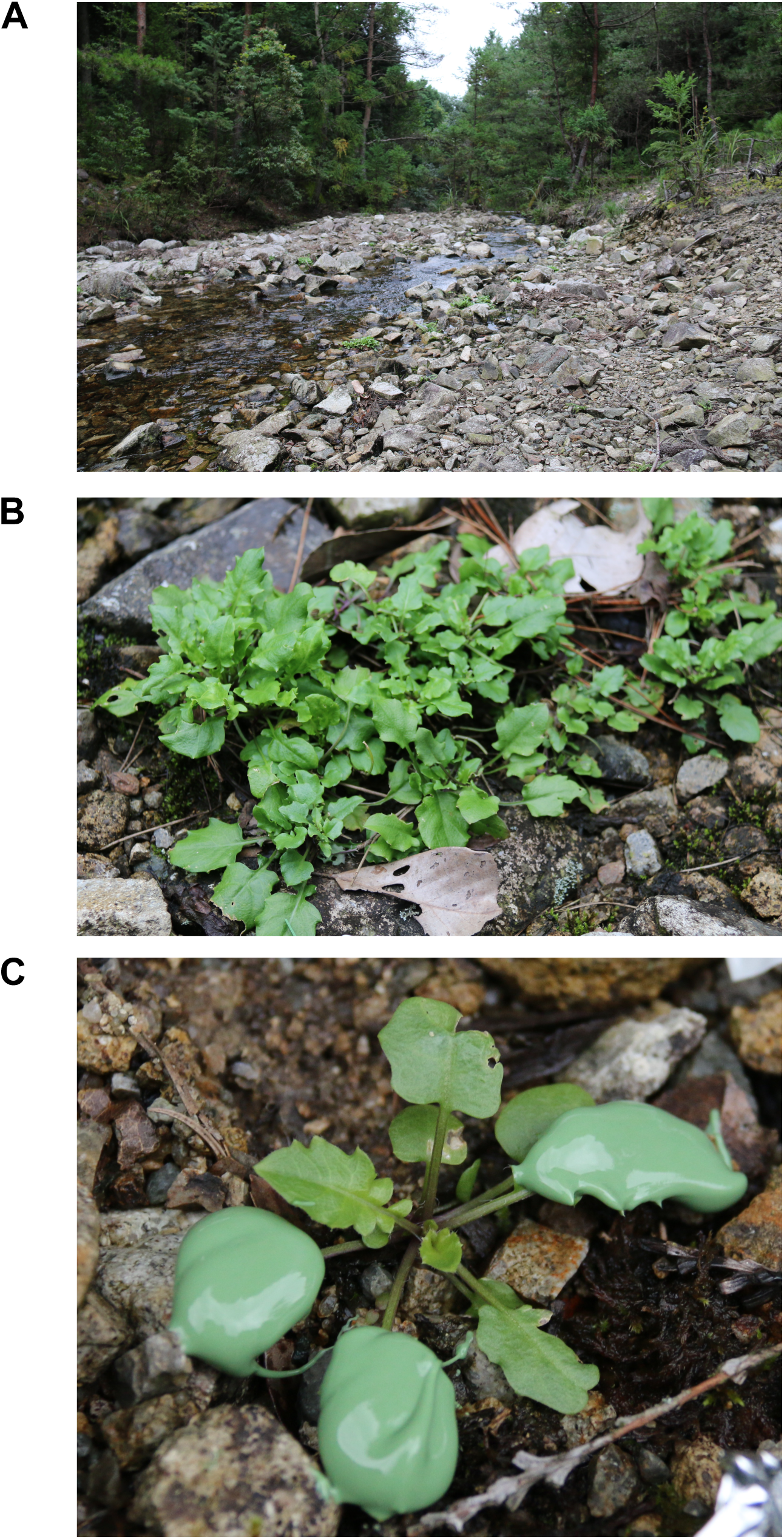
Field sampling of *Arabidopsis halleri* for stomatal density. (A) Overview of field site; (B) Rosette form of *A*. *halleri* plants when sampling during September; (C) Leaf surface impression acquisition using impression paste. The impression paste is green-coloured and occupies the surface of three rosette leaves.

### Stomatal density measurement

Eight plants of each trichome morph (hairy or glabrous) were selected at the study site, with individuals chosen such that the replicate plants were distributed evenly across the site. Glabrous and hairy morphs were identified by visual inspection of the leaf surface. It is thought that irradiance and ambient temperature are unlikely to influence the frequency of the morphs (Sato and Kudoh, 2017), but we cannot discount the possibility of microenvironment- or field site edge-effects. Stomatal density was measured by obtaining impressions from the adaxial surfaces of between 3 and 5 fully-expanded rosette leaves of each plant. We focused on the adaxial surface because this surface also harbours the majority of the trichomes. Between the times of 12:00 and 13:00, President Plus dental impression paste (Coltene) was applied to the adaxial side of each leaf to create a leaf surface impression (Fig. 1C). Solidified impression paste was removed from leaves and transported to the laboratory for further processing. First, each impression was assigned a randomly-generated number to ensure subsequent steps were performed blind. Each leaf impression was painted with transparent nail varnish (60 seconds super shine, Rimmel) that, after drying, was peeled away from the dental impression paste using transparent adhesive tape (Scotch Crystal). Next, the adhesive tape was used to attach the nail varnish impression to a 0.8 mm – 1 mm thick microscope slide. Leaf impressions were examined using an epifluorescence microscope in white light illumination mode. Images were captured from the centre of each leaf half, away from the midrib, using a Hamamatsu camera and Volocity software set to 20x zoom. Two images were captured from each impression, and the number of stomata and pavement cells was counted in an 800 µm x 800 µm square using the Fiji software to obtain cell density measures. Cell density measures were expressed as per mm^2^ (multiplication by 1.56).

In total, 29 and 31 leaf impressions were obtained in the field from hairy and glabrous plants, respectively. This produced 58 (hairy) and 62 (glabrous) microscopy images for analysis, because two images were captured from each impression. Stomatal index was calculated according to 1. After all measurements, data were disaggregated according to the blinding/randomization scheme. The differences between hairy and glabrous plants were statistically tested by nested analysis of variance, whereby the mean stomatal density or index per replicate plant was nested within the hairy and glabrous morphs. Tests were conducted using the R 3.6.0 software (Team, 2019) and plots generated with the beeswarm R package (v0.2.3) and Inkscape v0.91. No adjustments were applied to photographs in Fig. 1.

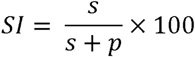

**Equation 1**. Derivation of stomatal index, where *SI* is stomatal index, *s* is the number of stomata in the field of view, and *p* is the number of epidermal pavement cells in the field of view.

## Results

We investigated stomatal patterning in naturally-occurring hairy and glabrous morphs of *A*. *halleri* (Sato and Kudoh, 2016). Approximately half of the *A*. *halleri* population at this study site is glabrous, whilst remaining plants have trichomes (Kawagoe et al., 2011). As trichome initiation occurs prior to stomatal meristemoid formation (Larkin et al., 1996; Glover, 2000), it is likely that trichome and stomatal patterning are linked (Bean et al., 2002), so we hypothesized that this might produce a difference in stomatal density between the two trichome morphs of *A*. *halleri* under natural conditions.

We found that the trichome formation dimorphism was accompanied by a difference in stomatal density (Fig. 2A; Fig. S1; Supplemental Dataset S1). Fully-expanded leaves of glabrous morphs had significantly greater stomatal density on the adaxial surface compared with hairy-leaved morphs (glabrous: 30.7 ± 2.8 stomata mm^-2^; hairy: 23.6 ± 2.3 stomata mm^-2^; mean ± s.e.m) (Fig. 2A; Table S1; Supplemental Dataset S1). Furthermore, the stomatal index of the adaxial surface was significantly greater in glabrous morphs (18.04 ± 0.92) compared with hairy morphs (16.44 ± 1.03) (Fig. 2B; Table S1). The adaxial surface pavement cell density did not differ significantly between the morphs (Table S1). Although leaf widths varied significantly among plants, they did not differ significantly between the morphs (glabrous: 12.4 ± 0.7 mm; hairy: 13.5 ± 1.0 mm; Fig. S2; Supplemental Dataset S1), suggesting that the stomatal density difference between the morphs is not due to differences in leaf expansion between the morphs (Table S1). Mean stomatal density ranged from 17 – 35 stomata mm^-2^ for hairy morphs and 24 – 49 stomata mm^-2^ for glabrous morphs (Fig. 2A). This stomatal density was lower than for *Arabidopsis thaliana*, which has reported stomatal densities of 180 – 350 stomata mm^-2^ depending on background accession and growth conditions (Gray et al., 2000; Zhang et al., 2008; Franks et al., 2015). Although lower than in *A*. *thaliana*, our measurements of stomatal density in *A*. *halleri* are consistent with a previous report of stomatal density of A. *halleri* subsp. *gemmifera*, which measured adaxial stomatal density of 46 stomata mm^-2^ at 430 m altitude in autumn rosette leaves, with stomatal density progressively increasing with greater altitudes (Aryal et al., 2018). Our field site was lower altitude (230 m), so the lower stomatal densities at our study site (Fig. 2A) are congruous with this previous study (Aryal et al., 2018).

**Figure 2.**
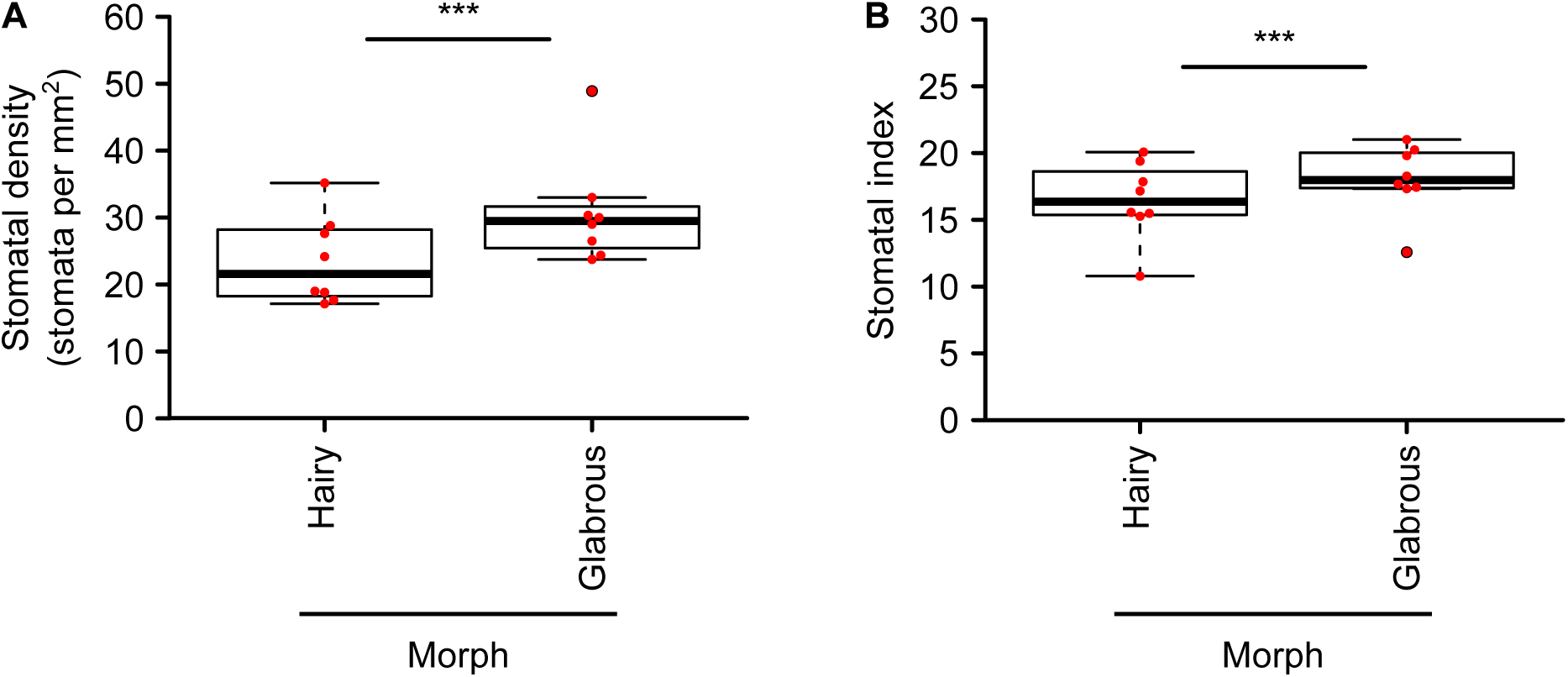
Stomatal density differs between hairy and glabrous morphs within a natural population of *Arabidopsis halleri*. (A) Stomatal density and (B) stomatal index for fully expanded leaves of hairy and glabrous morphs. Each red point represents the mean stomatal density or stomatal index from one individual plant. The mean stomatal density and stomatal index per plant was derived from 2 microscopy images analysed from 3-4 leaves of each plant. The centre line of the boxplot indicates the median. *n* = 8 plants from each morph; analysed by one-way nested ANOVA. *** indicates *p* < 0.001.

## Discussion

Glabrous plants had significantly greater stomatal density and stomatal index compared with hairy plants (Fig. 2A; Fig. 2B). As the density of surrounding pavement cells did not vary between the morphs, these differences in stomatal density and index are due to the greater density of stomata in glabrous morphs compared with hairy morphs (Fig. 2B). Our field data are consistent with a laboratory-based study in which transgenic tobacco plants expressing an *Antirrhinum myb*-like transcription factor, which caused an excess of trichomes, also had significantly reduced stomatal density (Glover et al., 1998). In a segregating tomato population, there is a negative correlation between stomatal and trichome density specifically under drought conditions (Galdon-Armero et al., 2018). Similarly, the trichome-bearing Col-0 accession of *A*. *thaliana* has lower stomatal density than the glabrous C24 accession (e.g. about 115 mm^-2^ for Col-0 and 180 mm^-2^ for C24) (Perazza et al., 1998; Lake and Woodward, 2008), although factors other than trichome density are likely to influence stomatal density between the accessions. This suggests that in natural populations of *A*. *halleri*, there could be a trade-off between trichome and stomatal development. Since the glabrous *gl1* mutant of *A*. *thaliana* has a significantly greater density of stomatal units compared with the wild type (Berger et al., 1998) and the glabrous phenotype of *A*. *halleri* at this study site is associated with an insertion within *GL1* (Kawagoe et al., 2011), it is possible that the *GL1* haplotype influences the stomatal density within this population of *A*. *halleri*.

In some cases, there does not appear to be a tradeoff between stomatal and trichome density. For example, elevated CO_2_ decreases stomatal density (Woodward and Kelly, 1995), but might also reduce trichome density (Bidart-Bouzat et al., 2005). Therefore, in future, it could be informative to examine the relationship between stomatal and trichome density under a range of different experimental conditions that apply different types of selection pressure. Furthermore, we sampled the adaxial leaf surface and it is possible that the presence of trichomes might affect stomatal density differently on the abaxial surface because, depending on environmental conditions, abaxial stomatal density of *A*. *halleri* can be 10—30% greater than the adaxial surface (Aryal et al., 2018).

Interestingly, trichome production appears to impose a fitness cost. For example, glabrous *A*. *halleri* plants have 10% greater biomass than hairy plants when grown in the absence of herbivores (Sato and Kudoh, 2016). This cost of herbivore resistance arising from trichome formation also occurs in glabrous and hairy *A*. *lyrata* (Løe et al., 2007; Sletvold et al., 2010) and *A*. *thaliana* (Mauricio and Rausher, 1997; Mauricio, 1998) under experimental conditions excluding herbivores. Whilst this fitness advantage of glabrous over hairy leaves in the absence of herbivory might be due to trichome production (Mauricio and Rausher, 1997; Mauricio, 1998; Kawagoe and Kudoh, 2010; Sletvold et al., 2010; Kawagoe et al., 2011; Sletvold and Ågren, 2012), we suggest that glabrous morphs might also gain an advantage by having a greater density or number of stomata. It has been proposed that increasing the number of stomata could increase carbon assimilation (Lawson and Blatt, 2014). For example, Arabidopsis overexpressing *STOMAGEN* has greater stomatal density and a 30% increase in carbon assimilation compared with the wild type. However, these lines also have higher transpiration rates and consequently lower water use efficiency (Tanaka et al., 2013). An alternative interpretation is that differences in the developmental program of the hairy and glabrous morphs might lead to differences in cell or leaf size, which ultimately causes the biomass difference between the morphs. In our samples, the width of fully expanded leaves did not differ significantly between the morphs (Fig. S2). Using these leaf width measures as a proxy for leaf size suggests that the biomass difference between the morphs is not due to leaf size differences between the morphs. However, there might be differences in other developmental characteristics that affect biomass, such as leaf thickness, which we did not compare between the morphs.

Optimal stomatal density is important to achieve high photosynthetic rates. A low stomatal density restricts CO_2_ vertical diffusion through the leaf and reduces photosynthetic rates, whilst high-density stomatal clustering diminishes CO_2_ diffusion and causes low carbon assimilation (Lawson and Blatt, 2014). Both *A*. *halleri* morphs examined are likely to be within an optimal range of stomatal densities, having evolved and survived under natural conditions. However, the higher stomatal density in the glabrous morph might contribute to its faster growth in absence of herbivory (Sato and Kudoh, 2016). In future, it would be interesting to explore this by measuring the CO_2_ assimilation rate of these trichome morphs under laboratory and/or natural conditions. It would also be informative to determine whether the stomatal density difference between the two trichome morphs confers any advantages within microenvironments characterized by differences in water or light availability. The lower stomatal density of *A*. *halleri* compared with *A*. *thaliana* (Gray et al., 2000; Zhang et al., 2008; Franks et al., 2015) might reflect differences in growth conditions. An alternative explanation might relate to genome size, because there appears to be a negative correlation between genome size and stomatal density (Beaulieu et al., 2008), and the genome of *A*. *halleri* (250 Mb) is approximately double the size of the *A*. *thaliana* genome (125 Mb) (The Arabidopsis Genome, 2000; Briskine et al., 2017).

In summary, we found that a glabrous morph of *A*. *halleri* growing under natural conditions had greater stomatal density and stomatal index than a hairy morph. This might contribute to the reported fitness advantage of glabrous plants over hairy plants in absence of herbivores (Sato and Kudoh, 2017). This differing stomatal density phenotype might derive from the common upstream components in the pathways leading to trichome and guard cell development.

## Acknowledgements

We thank Dora Cano-Ramirez, Haruki Nishio and Tasuku Ito for experimental assistance. This research was funded by the UK Biotechnology and Biological Sciences Research Council (BBSRC; SWBio DTP award BB/J014400/1 and Institute Strategic Programme GEN BB/P013511/1), The Royal Society (grant IE140501), and the Japan Society for Promotion of Science (JSPS; CREST no. JPMJCR15O1). This research was conducted through Joint Usage of the Center for Ecological Research, Kyoto University.

## Author contributions

NMLS, JS, MNH, SAT, GT, HK and AND performed experimentation and/or analysed data, and NMLS, MNH, HK and AND interpreted findings and wrote the paper.

## Data availability

All data generated during this study are included in the published article and Supplementary Information files.

## Figure legends

**Figure S1.** Stomatal density differs between hairy and glabrous morphs within a natural population of *Arabidopsis halleri*. Figure summarizes all data collected for (A) stomatal density and (B) stomatal index of fully expanded leaves of hairy and glabrous morphs. Each red point represents one measurement from one microscopy sample, with 2 samples analyzed from 3-4 leaves from 8 individual plants of each morph. The centre line of the boxplot indicates the median. Data represent a total of 58 and 62 microscopy images from hairy and glabrous plants, respectively.

**Figure S2.** Mean width of fully-expanded leaves does not differ between glabrous and hairy morphs within a natural population of *Arabidopsis halleri*. Each red point represents the mean leaf width from one plant, calculated from the width of either 3 or 4 leaves per plant. 8 replicate plants were sampled for each morph. The centre line of the boxplot indicates the median. Data were analyzed by one-way nested ANOVA, using the mean leaf width per plant as the level of replication. N. S. indicates difference is not statistically significant at *p* >= 0.05

**Table S1.** Nested ANOVA analysis of (a) stomatal density, (b) stomatal index, (c) pavement cell density and (d) leaf width. *Df*, degree of freedom; *, **, ***, significant at *p* < 0.05, < 0.01, < 0.001 respectively; NS, not significant at *p* >= 0.05.

**Dataset S1.** All stomatal density and leaf width data collected during experimentation.

## References

Aikawa S, Kobayashi MJ, Satake A, Shimizu KK, Kudoh H (2010) Robust control of the seasonal expression of the Arabidopsis FLC gene in a fluctuating environment. Proceedings of the National Academy of Sciences 107: 11632–11637

Aryal B, Shinohara W, Honjo MN, Kudoh H (2018) Genetic differentiation in cauline-leaf-specific wettability of a rosette-forming perennial Arabidopsis from two contrasting montane habitats. Annals of Botany 121: 1351–1360

Bean GJ, Marks MD, Hulskamp M, Clayton M, Croxdale JL (2002) Tissue patterning of Arabidopsis cotyledons. New Phytologist 153: 461–467

Beaulieu JM, Leitch IJ, Patel S, Pendharkar A, Knight CA (2008) Genome size is a strong predictor of cell size and stomatal density in angiosperms. New Phytologist 179: 975–986

Berger F, Linstead P, Dolan L, Haseloff J (1998) Stomata patterning on the hypocotyl of Arabidopsis thaliana is controlled by genes involved in the control of root epidermis patterning. Developmental Biology 194: 226–234

Bidart-Bouzat MG, Mithen R, Berenbaum MR (2005) Elevated CO^2^ influences herbivory-induced defense responses of Arabidopsis thaliana. Oecologia 145: 415–424

Bird SM, Gray JE (2003) Signals from the cuticle affect epidermal cell differentiation. New Phytologist 157: 9–23

Briskine RV, Paape T, Shimizu-Inatsugi R, Nishiyama T, Akama S, Sese J, Shimizu KK (2017) Genome assembly and annotation of Arabidopsis halleri, a model for heavy metal hyperaccumulation and evolutionary ecology. Molecular Ecology Resources 17: 1025–1036

Dalin P, Agren J, Bjorkman C, Huttunen P, Karkkainen K (2008) Leaf trichome formation and plant resistance to herbivory. In A Schaller, ed, Induced plant resistance to herbivory. Springer, pp 89–105

Franks PJ, W. Doheny-Adams T, Britton-Harper ZJ, Gray JE (2015) Increasing water-use efficiency directly through genetic manipulation of stomatal density. New Phytologist 207: 188–195

Galdon-Armero J, Fullana-Pericas M, Mulet PA, Conesa MA, Martin C, Galmes J (2018) The ratio of trichomes to stomata is associated with water use efficiency in Solanum lycopersicum (tomato). The Plant Journal 96: 607–619

Glover BJ (2000) Differentiation in plant epidermal cells. Journal of Experimental Botany 51: 497–505

Glover BJ, Perez-Rodriguez M, Martin C (1998) Development of several epidermal cell types can be specified by the same MYB-related plant transcription factor. Development 125: 3497

Gray JE, Holroyd GH, van der Lee FM, Bahrami AR, Sijmons PC, Woodward FI, Schuch W, Hetherington AM (2000) The HIC signalling pathway links CO^2^ perception to stomatal development. Nature 408: 713–716

Handley R, Ekbom B, Ågren J (2005) Variation in trichome density and resistance against a specialist insect herbivore in natural populations of Arabidopsis thaliana. Ecological Entomology 30: 284–292

Honjo MN, Kudoh H (2019) Arabidopsis halleri: a perennial model system for studying population differentiation and local adaptation. AoB PLANTS 11

Kawagoe T, Kudoh H (2010) Escape from floral herbivory by early flowering in Arabidopsis halleri subsp. gemmifera. Oecologia 164: 713–720

Kawagoe T, Shimizu KK, Kakutani T, Kudoh H (2011) Coexistence of trichome variation in a natural plant population: A combined study using ecological and candidate gene approaches. PLoS ONE 6: e22184

Kudoh H, Honjo MN, Nishio H, Sugisaka J (2018) The long-term “in natura” study sites of Arabidopsis halleri for plant transcription and epigenetic modification analyses in natural environments. In Plant Transcription Factors. Springer, pp 41–57

Lake JA, Woodward FI (2008) Response of stomatal numbers to CO^2^ and humidity: control by transpiration rate and abscisic acid. New Phytologist 179: 397–404

Larkin JC, Young N, Prigge M, Marks MD (1996) The control of trichome spacing and number in Arabidopsis. Development 122: 997–1005

Lawson T, Blatt MR (2014) Stomatal size, speed, and responsiveness impact on photosynthesis and water use efficiency. Plant Physiology 164: 1556–1570

Levin DA (1973) The role of trichomes in plant defense. The Quarterly Review of Biology 48: 3–15

Løe G, Toräng P, Gaudeul M, Ågren J (2007) Trichome production and spatiotemporal variation in herbivory in the perennial herb Arabidopsis lyrata. Oikos 116: 134–142

Mauricio R (1998) Costs of resistance to natural enemies in field populations of the annual plant Arabidopsis thaliana. The American Naturalist 151: 20–28

Mauricio R, Rausher MD (1997) Experimental manipulation of putative selective agents provides evidence for the role of natural enemies in the evolution of plant defense. Evolution 51: 1435–1444

Oppenheimer DG, Herman PL, Sivakumaran S, Esch J, Marks MD (1991) A myb gene required for leaf trichome differentiation in Arabidopsis is expressed in stipules. Cell 67: 483–493

Perazza D, Vachon G, Herzog M (1998) Gibberellins promote trichome formation by up-regulating GLABROUS1 in Arabidopsis. Plant Physiology 117: 375–383

R Core Team (2019) R: A language and environment for statistical computing. R Foundation for Statistical Computing, Vienna, Austria. URL https://www.R-project.org/.

Sato Y, Kudoh H (2015) Tests of associational defence provided by hairy plants for glabrous plants of Arabidopsis halleri subsp. gemmifera against insect herbivores. Ecological Entomology 40: 269–279

Sato Y, Kudoh H (2016) Associational effects against a leaf beetle mediate a minority advantage in defense and growth between hairy and glabrous plants. Evolutionary Ecology 30: 137–154

Sato Y, Kudoh H (2017) Fine-scale frequency differentiation along a herbivory gradient in the trichome dimorphism of a wild Arabidopsis. Ecology and Evolution 7: 2133–2141

Sletvold N, Ågren J (2012) Variation in tolerance to drought among Scandinavian populations of Arabidopsis lyrata. Evolutionary Ecology 26: 559–577

Sletvold N, Huttunen P, Handley R, Kärkkäinen K, Ågren J (2010) Cost of trichome production and resistance to a specialist insect herbivore in Arabidopsis lyrata. Evolutionary Ecology 24: 1307–1319

Tanaka Y, Sugano SS, Shimada T, Hara-Nishimura I (2013) Enhancement of leaf photosynthetic capacity through increased stomatal density in Arabidopsis. New Phytologist 198: 757–764

The Arabidopsis Genome Initiative (2000) Analysis of the genome sequence of the flowering plant Arabidopsis thaliana. Nature 408: 796–815

Woodward FI, Kelly CK (1995) The influence of CO^2^ concentration on stomatal density. New Phytologist 131: 311–327

Zhang L, Hu G, Cheng Y, Huang J (2008) Heterotrimeric G protein α and β subunits antagonistically modulate stomatal density in Arabidopsis thaliana. Developmental Biology 324: 68–75

