## Supplemental Table S1 and Supplemental Figure S1 and S2 for "Altered stomatal patterning accompanies a trichome dimorphism in a natural population of *Arabidopsis*"

**Table S1.** Nested ANOVA analysis of (a) stomatal density, (b) stomatal index, (c) pavement cell density and (d) leaf width. *Df*, degree of freedom;*, **, ***, significant at *p* < 0.05, < 0.01, < 0.001, respectively; NS, not significant at *p* >= 0.05.

*(a) Stomatal density*

| Factor | *Df* | Sum of square | Mean square | *F* value (divided by residuals) | *F* value (divided by plants) |
|---|---|---|---|---|---|
| Morph | 1 | 61.1 | 61.2 | 19.07*** | $2.118^{NS}$ |
| Plant (nested in morph) | 14 | 404 | 28.9 | 9.00*** | |
| Residuals | 44 | 141.1 | 3.21 | | |

*(b) Stomatal index*

| Factor | *Df* | Sum of square | Mean square | *F* value (divided by residuals) | *F* value (divided by plants) |
|---|---|---|---|---|---|
| Morph | 1 | 2177 | 2177 | 13.0*** | $3.994^{NS}$ |
| Plant (nested in morph) | 14 | 7631 | 545.1 | 3.26** | |
| Residuals | 44 | 7348 | 167.0 | | |

*(c) Pavement cell density*

| Factor | *Df* | Sum of square | Mean square | *F* value (divided by residuals) | *F* value (divided by plants) |
|---|---|---|---|---|---|
| Morph | 1 | 10070 | 10070 | $2.847^{NS}$ | $0.923^{NS}$ |
| Plant (nested in morph) | 14 | 152796 | 10914 | 3.086** | |
| Residuals | 44 | 155630 | 3537 | | |

*(d) Leaf width*

| Factor | *Df* | Sum of square | Mean square | *F* value (divided by residuals) | *F* value (divided by plants) |
|---|---|---|---|---|---|
| Morph | 1 | 12.35 | 12.35 | $2.77^{NS}$ | $0.59^{NS}$ |
| Plant (nested in morph) | 14 | 292.15 | 20.87 | $4.68^{***}$ | |
| Residuals | 44 | 196.34 | 4.46 | | |

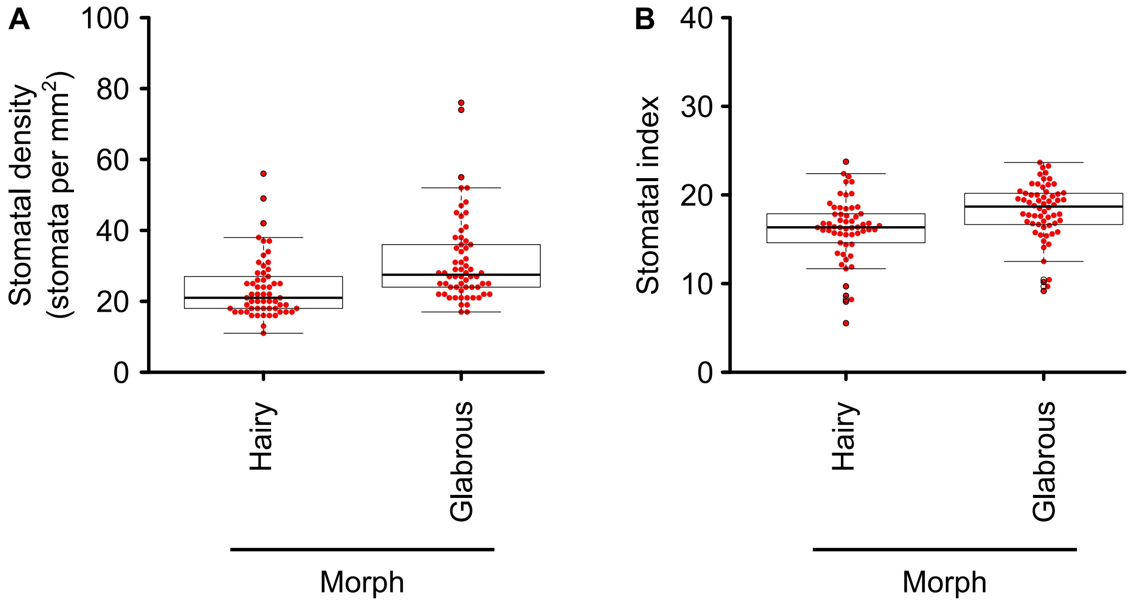


**Figure S1.** Stomatal density differs between hairy and glabrous morphs within a natural population of *Arabidopsis halleri*. Figure summarizes all data collected for (A) stomatal density and (B) stomatal index of fully expanded leaves of hairy and glabrous morphs. Each red point represents one measurement from one microscopy sample, with 2 samples analyzed from 3-4 leaves from 8 individual plants of each morph. The centre line of the boxplot indicates the median. Data represent a total of 58 and 62 microscopy samples from hairy and glabrous plants, respectively.

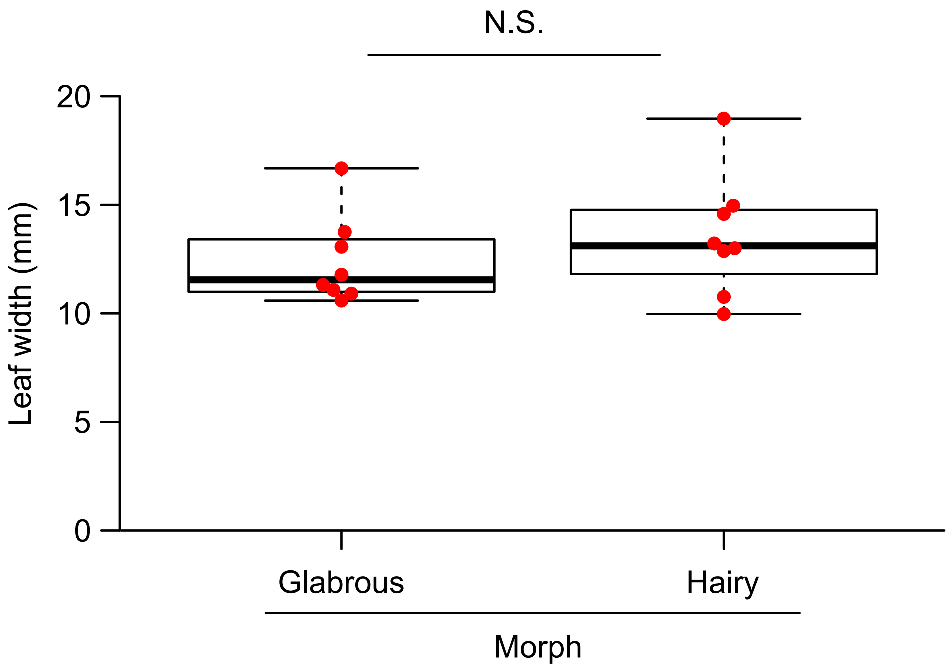


**Figure S2.** Mean width of fully-expanded leaves does not differ between glabrous and hairy morphs within a natural population of *Arabidopsis halleri*. Each red point represents the mean leaf width from one plant, calculated from the width of either 3 or 4 leaves per plant. 8 replicate plants were sampled for each morph. The centre line of the boxplot indicates the median. Data were analyzed by one-way nested ANOVA, using the mean leaf width per plant as the level of replication.
